# Outer membrane protein A (OmpA) deficient *Salmonella* Typhimurium displays enhanced susceptibility towards β-lactam antibiotics: third-generation cephalosporins (ceftazidime) and carbapenems (meropenem)

**DOI:** 10.1101/2022.01.16.476490

**Authors:** Atish Roy Chowdhury, Debapriya Mukherjee, Ashish Kumar Singh, Dipshikha Chakravortty

## Abstract

The invasive non-typhoidal serovar of *Salmonella enterica,* namely *Salmonella* Typhimurium ST313, causes bloodstream infection in sub-Saharan Africa. Like other bacterial pathogens, the development of antimicrobial resistance is a severe problem in curing non-typhoidal *Salmonella* infection. In this work, we have investigated the role of four prominent outer membrane porins of *S.* Typhimurium, namely OmpA, OmpC, OmpD, and OmpF, in resistance against broad-spectrum β-lactam antibiotics-ceftazidime and meropenem. We found that deleting OmpA from *Salmonella* makes the bacteria susceptible to β-lactam drugs. The MIC for both the antibiotics reduced significantly for STM *ΔompA* compared to the wild-type and the *ompA* complemented strains. Despite the presence of antibiotics, the uninterrupted growth of STM *ΔompC*, *ΔompD,* and *ΔompF* endorsed the dispensability of these three porins in antimicrobial resistance. The β-lactam antibiotics caused massive depolarization in the outer membrane of the bacteria in the absence of OmpA. We have proved that none of the extracellular loops but the complete structure of perfectly folded OmpA is required by the bacteria for developing antimicrobial resistance. Our data revealed that STM *ΔompA* consumed more antibiotics than the wild-type and the complemented strain, resulting in severe damage of the bacterial outer membrane and subsequent killing of the pathogen by antibiotic-mediated oxidative stress. Upon deleting *ompA*, the steady decrease in the relative proportion of antibiotic-resistant persisters and the clearance of the STM *ΔompA* from the liver and spleen of C57BL/6 mice upon treatment with ceftazidime proved the role of OmpA in rendering protection against β-lactam antibiotics.

## Introduction

*Salmonella enterica* is one of the leading causes of foodborne diseases and associated with infecting 10% of the population worldwide, with 33 million deaths annually [1]. *Salmonella* Typhimurium, the most commonly reported non-typhoidal serovar of this pathogen, causes self-limiting gastroenteritis in humans and typhoid fever-like symptoms in animals. Every year 1.3 billion cases of *Salmonella-*related gastroenteritis are reported globally, with approximately 3 million deaths [2]. *Salmonella* Typhimurium sequence types (ST) 19 and 34 are the primary reasons for global gastroenteritis [3]. However, bloodstream infection caused by the non-typhoidal *Salmonella* serovars is also a serious health hazard in sub-Saharan Africa (SSA) [4]. *S.* Typhimurium ST313, an invasive non-typhoidal serovar (iNTS) of *Salmonella,* causes severe bloodstream infection in malnourished children and HIV-infected adults in SSA and leads to innumerable deaths annually [5–7]. Over the past thirty years, the rapid emergence and subsequent global spread of multidrug-resistant *Salmonella* Typhimurium such as *S.* Typhimurium DT104 posed a severe threat to public health [8].

The outer membrane of Gram-negative bacteria is an asymmetrical lipid bilayer that consists of phospholipids in the inner leaflet and lipopolysaccharides in the outer leaflet. It acts as an impenetrable barrier and restricts the entry of many antimicrobials [9]. Apart from the lipopolysaccharides, the outer membrane of Gram-negative bacteria is densely populated with porins, an outer membrane-bound β barrel protein, which helps in the transportation of salts, sugars, nutrients, peptides, amino acids, vitamins, etc., across the impermeable outer membrane of bacteria [10]. Besides having a significant role in transport across the membrane, the porins maintain the integrity of the bacterial outer membrane. Moreover, Gram-negative bacteria change their outer membrane permeability using the porins to develop resistance against antibiotics, antimicrobial peptides, etc., [11, 12].

OmpA of *Escherichia coli* helps the bacteria to build up resistance against β-lactams, glycopeptides, amphenicol, and licosamides [12]. Conversely, OmpF facilitates the transportation of β lactam antibiotics across the outer membrane, making the bacteria susceptible to antibiotic treatment [12]. OmpA is responsible for the multidrug resistance (MDR) phenotype in *Acinetobacter baumannii* by providing resistance against nalidixic acid, chloramphenicol, and aztreonam [13]. The deletion of the OmpA-like domain (amino acids 223-356) from the structure of OmpA increases the susceptibility of *Acinetobacter baumannii* towards imipenem, gentamycin, trimethoprim, and aztreonam, suggesting the mechanistic insight into the drug-resistance of *Acinetobacter* [14]. On the contrary, it has also been reported that the OmpA in *Acinetobacter baumannii* acts as a selective porin, mediating the passage of ETX_2514_, a β lactamase inhibitor, and further enhances the antibacterial activity of sulbactam [15]. The deletion of OmpA from *Klebsiella pneumoniae* enhances the susceptibility of the bacteria towards antimicrobial peptides such as polymixin B and protamine [16].

OmpA, OmpC, OmpD, and OmpF are the most abundant porins found on the outer membrane of *Salmonella* Typhimurium [17]. Earlier, we reported that OmpA protects the intracellular *Salmonella* from nitrosative stress in murine macrophages. The deletion of *ompA* from *Salmonella* resulted in the overexpression of *ompC*, *ompD,* and *ompF*. The same study revealed that the enhanced expression of OmpF in *Salmonella* lacking OmpA makes the bacteria susceptible to *in vitro* and *in vivo* nitrosative stress [18]. However, the role of these *Salmonella* porins (OmpA, OmpC, OmpD, and OmpF) in antibiotic resistance is yet to be explored. In the current study, we have investigated the contribution of these porins in promoting bacterial resistance against two β lactam antibiotics, namely ceftazidime- a third-generation cephalosporin and meropenem- a carbapenem drug. Both antibiotics can inhibit the growth of bacteria by interfering with the cell wall biosynthesis after binding to penicillin-binding proteins [19, 20]. Our data revealed that out of all four porins, only OmpA provides *S.* Typhimurium with a substantial amount of protection against ceftazidime and meropenem. We found that the externally exposed extracellular loops of OmpA have a very feeble role in maintaining anti-microbial resistance. Instead, deleting OmpA from *Salmonella* Typhimurium facilitated the entry of β-lactam antibiotics into the bacteria and caused a massive disruption in the outer membrane. To best our knowledge, this is the first study reporting the protective role of *S.* Typhimurium OmpA against broad-spectrum β lactam antibiotics.

## Results

### Deleting OmpA from *Salmonella* Typhimurium reduces the MIC for β-lactam antibiotics

Multiple studies have provided substantial evidence on the contributions of porins to maintain the outer membrane stability of *Salmonella* Typhimurium during *in vitro* and *in vivo* oxidative and nitrosative stresses [17, 18]. However, the precise role of *S.* Typhimurium outer membrane porins, namely OmpA, OmpC, OmpD, and OmpF, in antimicrobial resistance is yet to be tested. We tested the sensitivity of STM (WT), *ΔompA*, *ΔompC*, *ΔompD*, and *ΔompF* against two broad-spectrum β lactam antibiotics, namely ceftazidime and meropenem [19, 20]. Our data revealed that the deletion of OmpA from *Salmonella* has a tremendous impact on the ability of the bacteria to survive during antibiotic stress.

The enhanced sensitivity of the *ompA* knockout bacteria in the presence of ceftazidime and meropenem suggested a protective role of OmpA against the cell wall biosynthesis inhibitors **(Figure 1A and 1B)**. Surprisingly the deletion of other major porins of bacterial outer membrane- OmpC, OmpD, and OmpF did not show any significant growth inhibition when incubated with increasing concentrations of ceftazidime **(Figure 1A)**and meropenem **(Figure 1B)**. Resazurin assay was performed to validate this observation, which helped us estimate the viability of the bacteria in the presence of ceftazidime **(Figure S1A and 1B)** and meropenem **(Figure S2A and S2B)**. The percent viability of the OmpA deficient *Salmonella* was significantly reduced (49.8%) compared to the wild-type *Salmonella* (80.8%) in the presence of 0.5 μg/ mL concentration of ceftazidime **(Figure S1A and S1B)**. Likewise, when the bacteria were treated with 0.125 μg/ mL concentration of meropenem, we observed a severely compromised viable population of STM *ΔompA* (37.5%) compared to the STM (WT) (88.4%) **(Figure S2A and S2B)**. The MIC of ceftazidime and meropenem for wild-type *Salmonella spp.* are 1 and 2 μg/ mL, respectively [21]. Altogether, our data suggested that the deletion of OmpA from *Salmonella* Typhimurium reduced its MIC for ceftazidime to 0.5 μg/ mL and meropenem to 0.125 μg/ mL. On the contrary, the *ompC*, *ompD*, and *ompF* knockout strains exhibited an uninterrupted growth in increasing concentrations of both antibiotics, which endorsed the dispensability of these porins in developing antimicrobial resistance in *Salmonella* Typhimurium. While incubating with the sub-lethal concentrations of ceftazidime and meropenem, the partial recovery of the bacterial growth in *ompA* complemented strain of *Salmonella* strongly supported our previous observations **(Figure 1C and 1D)**. These results helped us conclude that, indeed, outer membrane protein A (OmpA) is required by *Salmonella* Typhimurium to build up resistance against β-lactam antibiotics.

**Figure 1.**
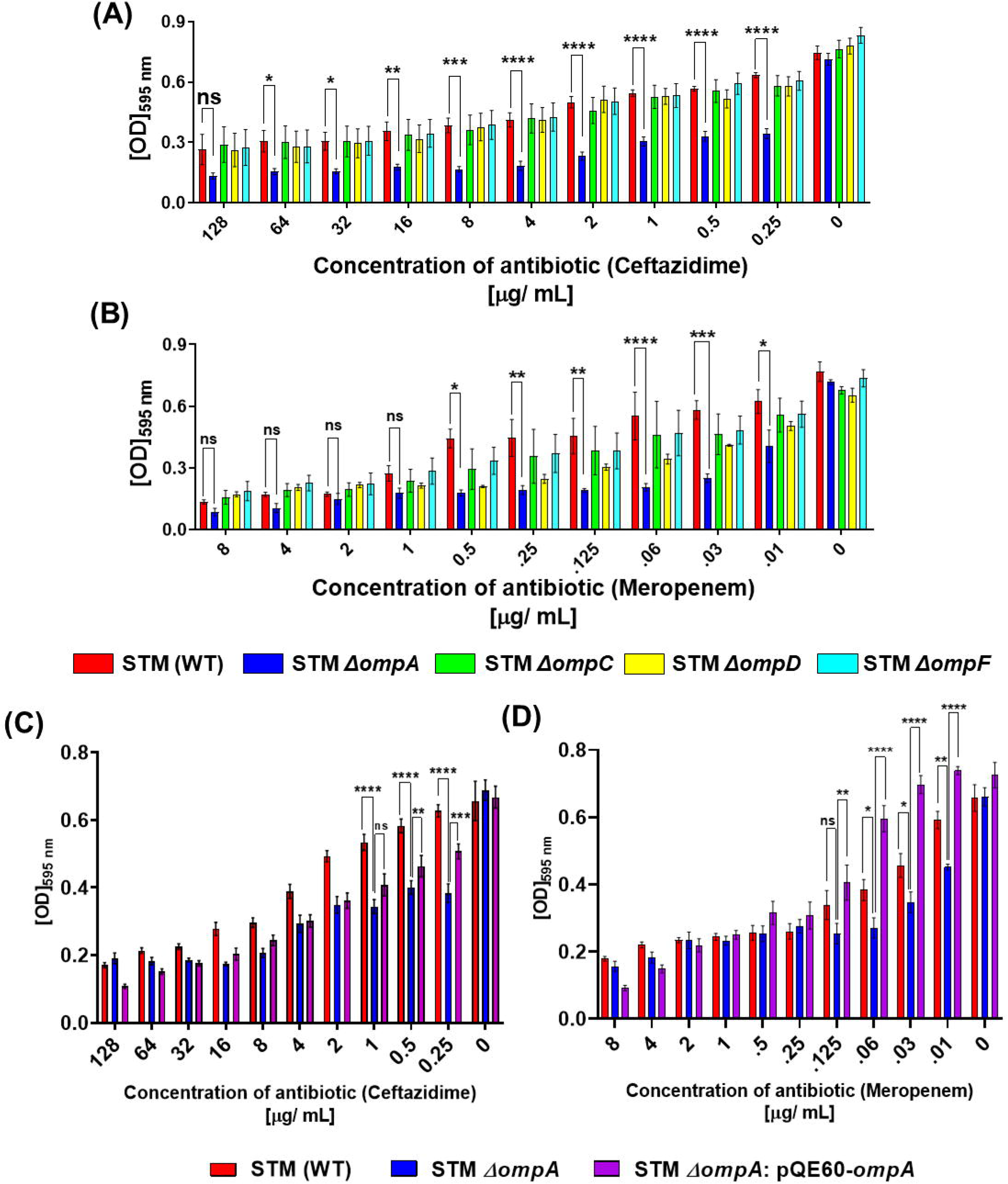
Deletion of OmpA from *Salmonella* Typhimurium reduced the minimal inhibitory concentration (MIC) for β-lactam antibiotics. Determination of minimal inhibitory concentration (MIC) for β-lactam antibiotics (A) ceftazidime (N=7) and (B) meropenem (N=4) for STM (WT), *ΔompA*, *ΔompC*, *ΔompD*, and *ΔompF* growing in cation-adjusted Mueller-Hinton broth. Studying the growth inhibition of STM (WT), *ΔompA*, and *ΔompA*:pQE60*-ompA*, growing in cation-adjusted Mueller-Hinton broth in the presence of varying concentrations of (C) ceftazidime (N=6) and (D) meropenem (N=4). ***(P)* *< 0.05, *(P)* **< 0.005, *(P)* ***< 0.0005, *(P)* ****< 0.0001, ns= non-significant, (2way ANOVA)**.

### The β-lactam antibiotics enhanced depolarization of the bacterial outer membrane in the absence of OmpA

Antimicrobial peptides kill bacterial pathogens by causing depolarization of the bacterial outer membrane [22–25]. Antibiotics such as ramoplanin, a peptidoglycan biosynthesis of the Gram-positive bacteria, can cause membrane depolarization in *Staphylococcus aureus* [26]. We hypothesized that the enhanced antibiotic-dependent killing of STM *ΔompA* is because of the greater depolarization of the bacterial outer membrane. A dye named DiBAC_4_ was used to measure the outer membrane depolarization of the bacteria treated with antibiotics. When the bacterial outer membrane is depolarized, the negative charge density of the bacterial cytoplasm reduces, which facilitates the entry and accumulation of DiBAC_4_ into the cell. To test our hypothesis, STM (WT), *ΔompA,* and *ΔompA*: pQE60- *ompA*, incubated with the increasing concentrations of β-lactam drugs, were treated with DiBAC_4_. The extent of membrane depolarization was measured by quantifying the DiBAC_4_ positive population with flow cytometry **(Figure 2)**. Deleting OmpA from *Salmonella* did not depolarize the outer membrane without antibiotics **(Figure 2A.II and 2B.II)**. But when the *ompA* knockout bacterial cells were incubated with three different concentrations of ceftazidime **(Figure 2A.III, 2A.IV, 2A.V, and 2A.VI)**, there was massive induction of outer membrane depolarization **(Figure 2A.III**-36.33%, **2A.IV-**47.18%, **2A.V-**53.96%, and **2A.VI-** cumulative trend) compared to the wild-type **(Figure 2A.III-**20.74%, **2A.IV-**31.19%, **2A.V-**29.31%, and **2A.VI-** cumulative trend) and the complemented strains **(Figure 2A.III-**15.4%, **2A.IV-**30.83%, and **2A.V-**35.34%, and **2A.VI-**cumulative trend). To see whether a similar kind of effect was exerted by meropenem, STM (WT), *ΔompA,* and *ΔompA*: pQE60-*ompA* were treated with 0.01, 0.03, and 0.06 μg/ mL concentrations of meropenem **(Figure 2B)**. In line with our expectations, an elevated depolarization in the outer membrane of STM *ΔompA* **(Figure 2B.III-**35.93%, **2B.IV-**44.08%, **and 2B.V-**49.83%, and **2B.VI-**cumulative trend) than STM (WT) **(Figure 2B.III-**30.62%, **2B.IV-**37.58% **and 2B.V-**36.06%, and **2B.VI-**cumulative trend) upon meropenem treatment was observed. The complementation of *ompA* in knockout bacteria efficiently reversed the depolarization phenotype **(Figure 2B.III-**8.26%, **2B.IV-**8.48%, and **2B.V-**22.07%, and **2B.VI-**cumulative trend). With an increase in the sub-lethal concentrations of β lactam drugs, the consistent elevation in the DiBAC_4_ positive population of STM *ΔompA* compared to the wild-type and complemented strains suggested that the lack of OmpA enhances the outer membrane permeability of the bacteria in response to antibiotics.

**Figure 2.**
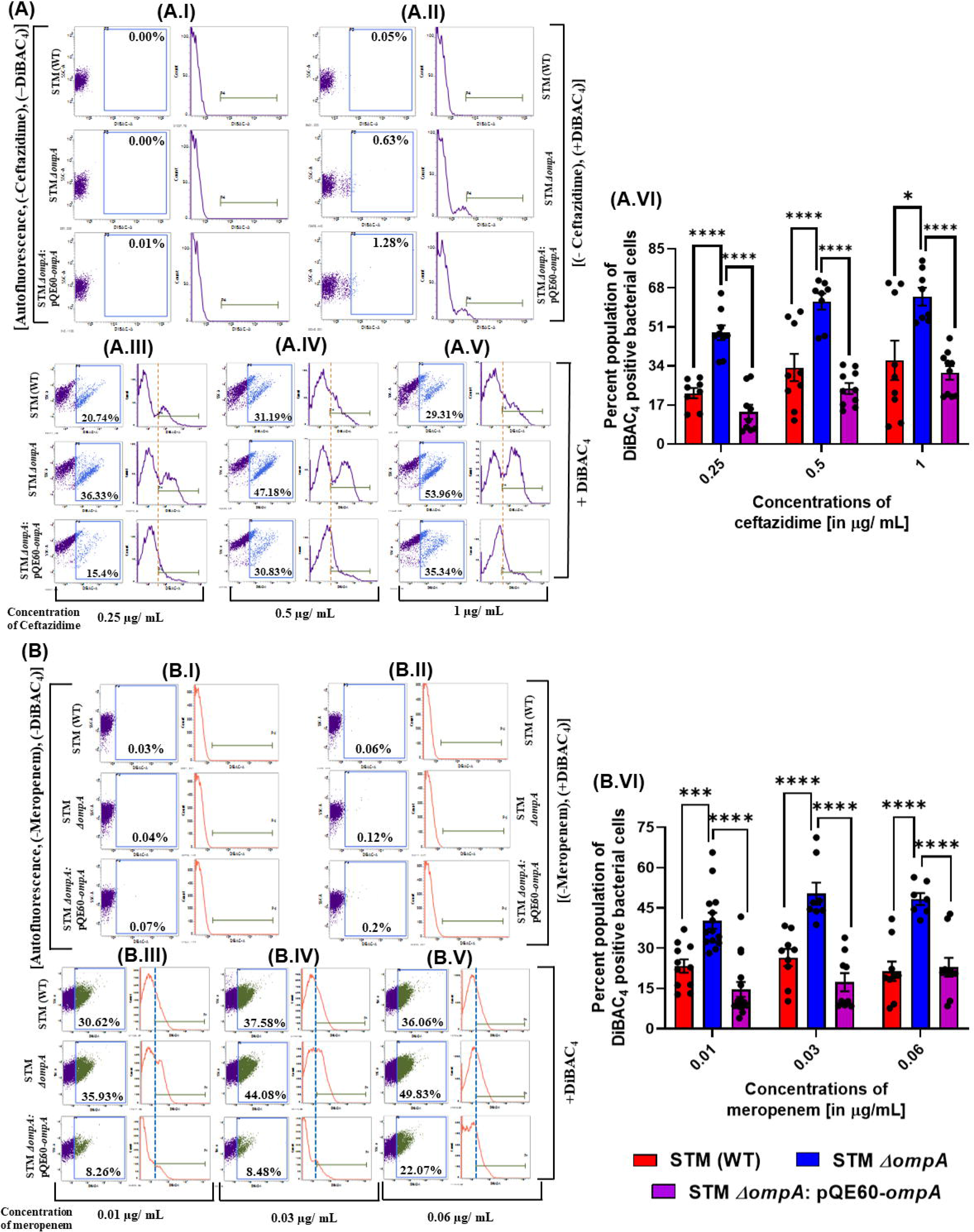
STM *ΔompA* showed enhanced outer membrane depolarization compared to the wild-type and complemented strain in the presence of β-lactam antibiotics. Measuring the outer membrane depolarization of STM (WT), *ΔompA*, and *ΔompA*:pQE60*-ompA*, growing in cation-adjusted Mueller-Hinton broth in the presence of increasing concentrations of (A) ceftazidime (A.I-autofluorescence, A.II-no-antibiotic control, A.III- 0.25, A.IV- 0.5, A.V- 1 μg/ mL, and A.VI- cumulative trend, n=2, N=5) and (B) meropenem (B.I- autofluorescence, B.II- no-antibiotic control, B.III- 0.01, B.IV- 0.03, and B.V- 0.06 μg/ mL, B.VI- cumulative trend, n=2, N=7) by DiBAC_4_ staining by flow cytometry. The final concentration of DiBAC_4_ used to measure the membrane depolarization was 1 μg/ mL. The representative image corresponds to one single experiment from the independently done experiments. The dot plot (SSC-A Vs. DCFDA-A) and the histogram (Count Vs. DCFDA-A) have been obtained from BD FACSuite software. ***(P)* *< 0.05, *(P)* **< 0.005, *(P)* ***< 0.0005, *(P)* ****< 0.0001, ns= non-significant, (2way ANOVA)**.

### None of the extracellular loops but the complete structure of OmpA shields the bacteria from antibiotic-mediated outer membrane depolarization

The β sheets of OmpA are connected to each other by four externally exposed extracellular loops. Earlier, we have reported that introducing mutations in these extracellular loops doesn’t alter the folding, expression, and outer membrane localization of OmpA in *Salmonella* Typhimurium [27]. We hypothesized that these extracellular loops could compensate for the function of whole OmpA in developing antibiotic resistance in *Salmonella*. By site-directed mutagenesis, multiple mutations were introduced to the loops of OmpA, and the loop mutants (STM *ΔompA*:pQE60-*ompA*-L1-1, *ΔompA*:pQE60-*ompA*-L1-2, *ΔompA*:pQE60-*ompA*-L2-1, *ΔompA*:pQE60-*ompA*-L2-2, *ΔompA*:pQE60-*ompA*-L3-1, and *ΔompA*:pQE60-*ompA*-L4-1) were subsequently subjected to antibiotic treatment **(Figure 3A and 3B)**. Surprisingly, it was observed that tampering with the loops didn’t exhibit any considerable impact on the survival of the bacteria in the presence of antibiotics **(Figure 3A and 3B)**. As we have observed earlier, the growth of STM *ΔompA* was inhibited at lower concentrations (<MIC) of ceftazidime **(Figure 3A)** and meropenem **(Figure 3B)** compared to the wild-type and the complemented strain. However, compared to STM *ΔompA,* the better survival of the OmpA extracellular loop mutants in the presence of β-lactam drugs proved that none of these extracellular loops could compensate for the role of perfectly folded whole OmpA in defending the bacteria from antibiotics. To validate this observation, the outer membrane depolarization of the *ompA* deficient *Salmonella* was measured along with the wild-type, complemented, and loop mutant strains by DiBAC_4_ staining **(Figure 3C, 3D, and 3E)**. Compared to the wild-type (ceftazidime- 11.66% and meropenem-26.89%) and the complemented (ceftazidime-4.07% and meropenem- 30.67%) strains, STM *ΔompA* showed enhanced outer membrane depolarization (ceftazidime- 17.76% and meropenem-38.68%) upon ceftazidime and meropenem treatment, which was significantly reduced in the OmpA loop mutants (ceftazidime-STM *ΔompA*:pQE60-*ompA*-L1-1- 3.93%, *ΔompA*:pQE60-*ompA*-L1-2-4.31%, *ΔompA*:pQE60-*ompA*-L2-1- 6.22%, *ΔompA*:pQE60-*ompA*-L2-2- 6.64%, *ΔompA*:pQE60-*ompA*-L3-1- 7.44%, & *ΔompA*:pQE60-*ompA*-L4-1- 11.27% and meropenem-STM *ΔompA*:pQE60-*ompA*-L1-1- 25.9%, *ΔompA*:pQE60-*ompA*-L1-2- 27.39%, *ΔompA*:pQE60-*ompA*-L2-1-12.56%, *ΔompA*:pQE60- *ompA*-L2-2-0.26%, *ΔompA*:pQE60-*ompA*-L3-1-23.7%, & *ΔompA*:pQE60-*ompA*-L4-1-3.66%) **(Figure 3C, 3D and 3E).** This result was further corroborated by estimating the percent viability of the wild-type, knockout, complemented, and OmpA loop mutant strains under ceftazidime treatment by resazurin assay **(Figure 3F)**. The better survival of the wild-type, complemented and loop mutant strains compared to STM *ΔompA* after ceftazidime treatment strongly suggested that the complete OmpA is required for antimicrobial resistance in *Salmonella* Typhimurium.

**Figure 3.**
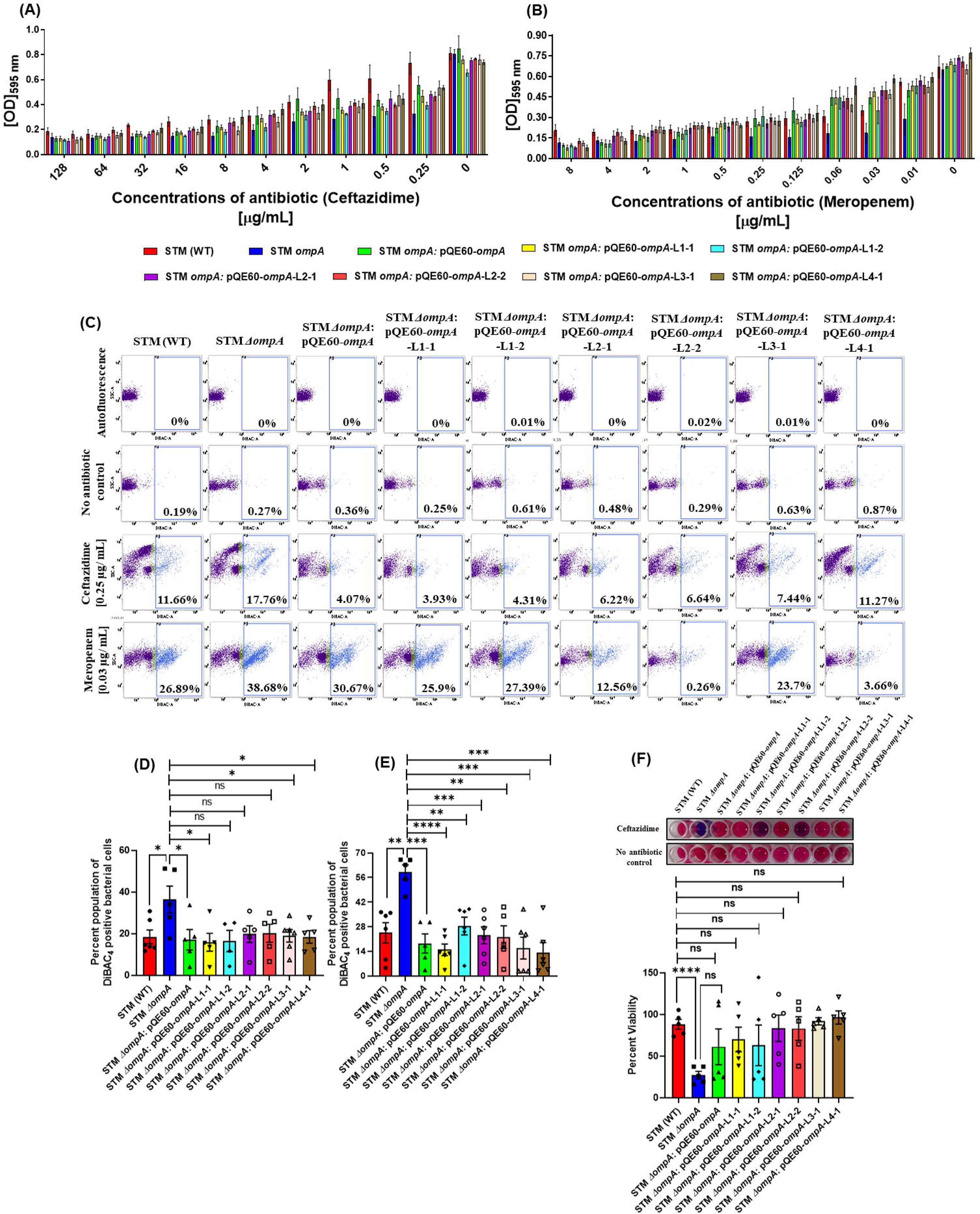
None of the extracellular loops but the complete OmpA protects *Salmonella* from β- lactam drugs. Examining the inhibition of the *in vitro* growth of STM (WT), *ΔompA*, *ΔompA*:pQE60-*ompA*, *ΔompA*:pQE60-*ompA*-L1-1, *ΔompA*:pQE60-*ompA*-L1-2, *ΔompA*:pQE60-*ompA*-L2-1, *ΔompA*:pQE60-*ompA*-L2-2, *ΔompA*:pQE60-*ompA*-L3-1, and *ΔompA*:pQE60-*ompA*-L4-1 growing in MH broth in the presence of (A) ceftazidime (N=5) and (B) meropenem (N=5). (C) The estimation of the outer membrane depolarization of STM (WT), *ΔompA*, *ΔompA*:pQE60- *ompA*, *ΔompA*:pQE60-*ompA*-L1-1, *ΔompA*:pQE60-*ompA*-L1-2, *ΔompA*:pQE60-*ompA*-L2-1, *ΔompA*:pQE60-*ompA*-L2-2, *ΔompA*:pQE60-*ompA*-L3-1, and *ΔompA*:pQE60-*ompA*-L4-1 in the presence of ceftazidime (0.25 μg/ mL) and meropenem (0.03 μg/ mL) by flowcytometry (n=2, N=3). The dot plot (SSC-A Vs. DCFDA-A) and the histogram (Count Vs. DCFDA-A) have been obtained from BD FACSuite software. The graphical representation of DiBAC_4_ positive population for (D) ceftazidime and (E) meropenem. The measurement of the percent viability of STM (WT), *ΔompA*, *ΔompA*:pQE60-*ompA*, *ΔompA*:pQE60-*ompA*-L1-1, *ΔompA*:pQE60-*ompA*-L1-2, *ΔompA*:pQE60-*ompA*-L2-1, *ΔompA*:pQE60-*ompA*-L2-2, *ΔompA*:pQE60-*ompA*-L3-1, and *ΔompA*:pQE60-*ompA*-L4-1 upon ceftazidime treatment by resazurin assay (n=2, N=3). ***(P)* *< 0.05, *(P)* **< 0.005, *(P)* ***< 0.0005, *(P)* ****< 0.0001, ns= non-significant, (unpaired student’s t-test)**.

### The enhanced uptake of β-lactam antibiotics by STM *ΔompA* caused severe damage to the bacterial outer membrane and made the bacteria susceptible to ROS

We further hypothesized that the enhanced uptake of β-lactam antibiotics by STM *ΔompA* results in the damage of the bacterial outer membrane, which eventually induces the depolarization of the bacterial outer membrane. To prove the increased consumption of antibiotics, the log-phase culture of STM (WT), *ΔompA*, and *ΔompA*:pQE60-*ompA* were treated with a very high concentration of meropenem (100-150 μg/ mL) for an hour and the remaining concentration of meropenem in the culture was quantified by HPLC to estimate the antibiotic consumption by bacteria **(Figure 5A and 5B)**. In line with our expectation, we have found that the remaining concentration of meropenem for STM *ΔompA* containing media was significantly lower than the wild-type bacteria, suggesting a higher intake of antibiotics by the mutant bacteria **(Figure 5A and 5B)**. To show the membrane disruption of *Salmonella* upon antibiotic treatment, all three bacterial strains were subjected to the increasing concentration of meropenem (0, 0.01, 0.03, and 0.06 μg/ mL) **(Figure 5C)**. Our data revealed that in the absence of antibiotics, the bacterial DNA (green) was tightly enclosed by an intact outer membrane **(Figure 5C)**. The increasing concentration of meropenem caused massive damage to the outer membrane of STM *ΔompA* compared to the wild-type and the complemented strains **(Figure 5C)**, which ultimately resulted in the release of bacterial DNA, followed by the death of the bacteria. To validate this observation, we assessed the morphology of the bacteria treated with 0.03 μg/ mL concentration of meropenem by atomic force microscopy **(Figure 5D)**. In continuation with the previous observation, we have found that the meropenem treatment severely impaired the morphology of STM *ΔompA* compared to the wild-type and the complemented strain **(Figure 5D)**. Altogether, our data suggest that the β-lactam antibiotics can induce immense disruption of the outer membrane of *Salmonella* Typhimurium in the absence of OmpA. Irrespective of their mode of action, most bactericidal antibiotics induce oxidative stress to kill bacterial pathogens [28]. The produced ROS can oxidize bacterial genomic DNA, membrane lipids, cellular proteins, etc. [29]. Both ceftazidime and meropenem can produce ROS while inhibiting bacterial growth. Apart from binding to the penicillin-binding proteins of rapidly dividing bacterial cells, ceftazidime can cause the oxidation of bacterial membrane lipids and the DNA bases [30]. The ROS-inducing ability of meropenem has also been tested in the case of another Gram-negative pathogen, *Burkholderia cepacia* and *Escherichia coli* [31, 32]. We have performed DCFDA staining of the bacteria and quantified the generation of intracellular ROS upon antibiotic treatment by flow cytometry. Treating the wild-type and *ompA* knockout bacteria with sub-lethal concentrations of ceftazidime (0.25 μg/ mL) and meropenem (0.01 μg/ mL) produced a comparable amount of ROS. Upon ceftazidime treatment for 18 to 24 hours, 64.33% of the STM (WT) and 64.26% of the STM *ΔompA* produced ROS **(Figure S3A.I, S3A.II, and S3A.III)**. At the same time, 12.8% of the wild-type and 11.29% of the *ompA* knockout bacteria had ROS when they were incubated with meropenem **(Figure S3B.I, S3B.II, and S3B.III)**. We further concluded that irrespective of the presence or absence of OmpA, the bacteria experience equivalent amount of oxidative stress upon antibiotic treatment. However, the higher outer membrane depolarization in the absence of OmpA makes the bacteria highly susceptible to antibiotic-dependent oxidative damage.

**Figure 4.**
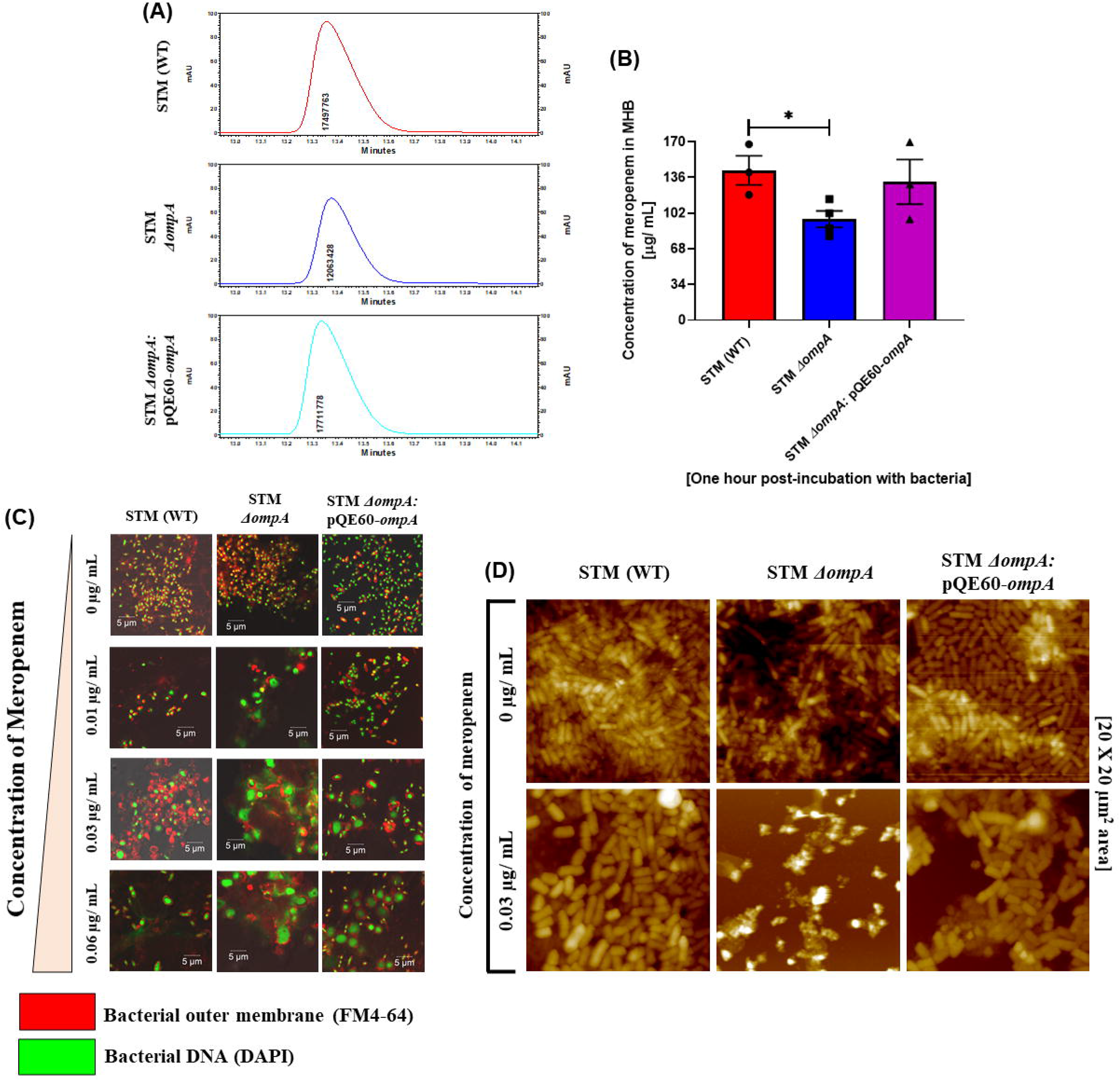
β-lactam antibiotics-dependent damage of the bacterial outer membrane. The (A) graphical representation and (B) the quantification of the amount of antibiotics entering the log phase cultures of STM (WT), *ΔompA*, *ΔompA*:pQE60-*ompA* growing for an hour in Muller-Hinton broth (N=4). (C) Pictorial representation of the outer membrane damage of STM (WT), *ΔompA*, and *ΔompA*:pQE60*-ompA*, growing in the presence of increasing concentrations of meropenem (0, 0.01, 0.03 and 0.06 μg/ mL, respectively) (N=3). The outer membrane and the DNA of the bacteria were stained with FM 4-64 (red) and DAPI (green- pseudo colour), respectively. The representative image represents one single experiment of three independently done experiments. **(Scale bar= 5 μm)**. (D) Atomic force micrograph to study the bacterial morphology in the presence and absence of meropenem. A 20X20 μm^2^ area from each coverslip having dried bacterial samples (antibiotic treated or untreated) were used for image acquisition. ***(P)* *< 0.05, ns= non-significant, (unpaired student’s t-test)**.

**Figure 5.**
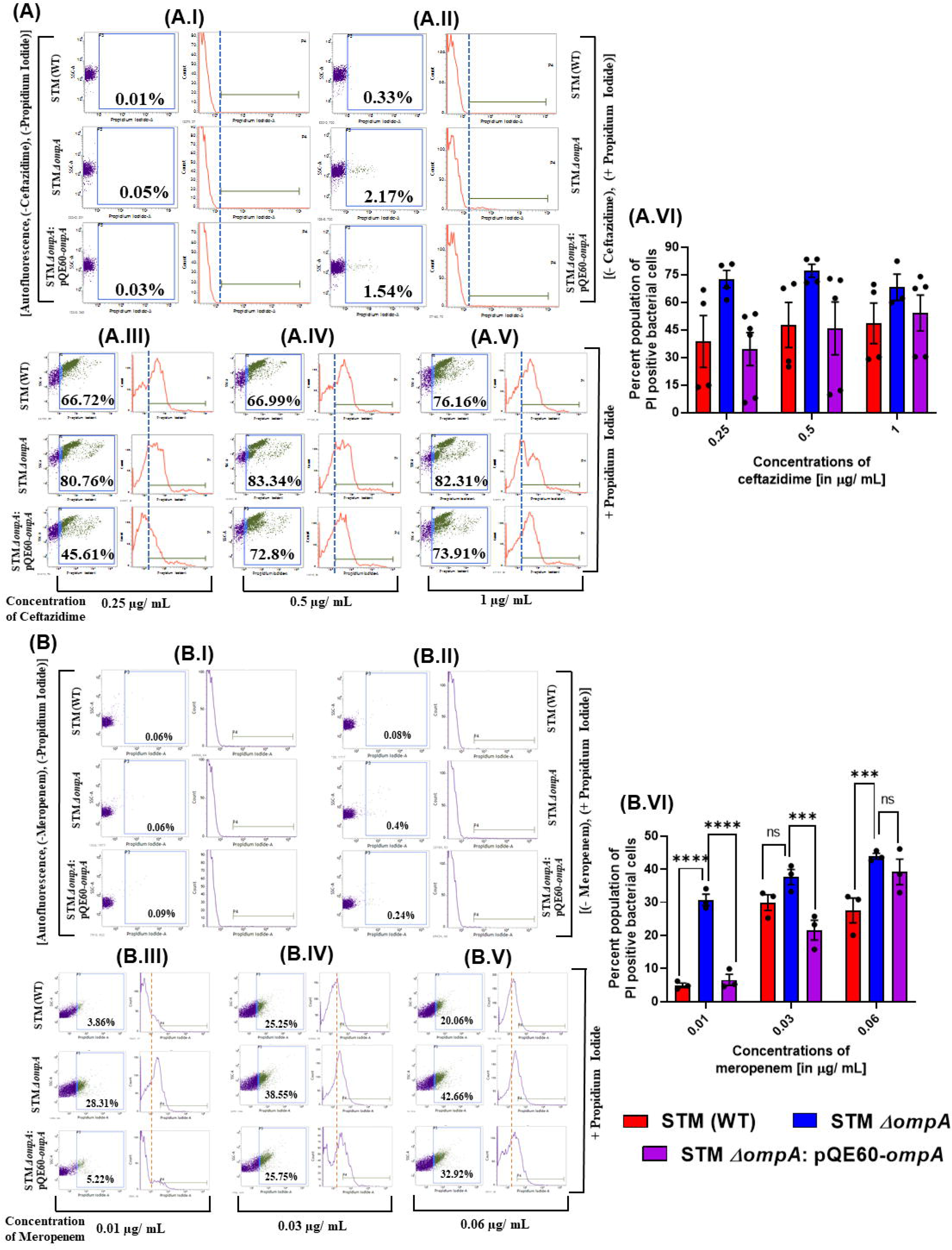
After β-lactam antibiotic treatment, the cell death indeced in STM *ΔompA* was more compared to the wild-type and the complemented strain. Estimating the death of STM (WT), *ΔompA*, and *ΔompA*:pQE60*-ompA*, growing in cation- adjusted Mueller-Hinton broth in the presence of increasing concentrations of (A) ceftazidime (A.I- autofluorescence, A.II- no-antibiotic control, A.III- 0.25, A.IV- 0.5, A.V- 1 μg/ mL, and A.VI- cumulative trend, n=2, N=3) and (B) meropenem (B.I- autofluorescence, B.II- no- antibiotic control, B.III- 0.01, B.IV- 0.03, and B.V- 0.06 μg/ mL, B.VI- cumulative trend, N=3) by propidium iodide staining by flow cytometry. The final concentration of propidium iodide used to measure the membrane percent cell death was 1 μg/ mL. The representative image corresponds to one single experiment from the independently done experiments. The dot plot (SSC-A Vs. DCFDA-A) and the histogram (Count Vs. DCFDA-A) have been obtained from BD FACSuite software. ***(P)* *< 0.05, *(P)* **< 0.005, *(P)* ***< 0.0005, *(P)* ****< 0.0001, ns= non-significant, (2way ANOVA)**.

### The absence of OmpA in *Salmonella* Typhimurium results in the enhanced killing of the bacteria in the presence of β-lactam antibiotics

We further wanted to correlate the impact of antibiotic-mediated membrane depolarization and subsequent oxidative damage with the survival of the bacteria. To investigate bacterial survival, we have stained the bacteria with propidium iodide (PI), which can cross the dead bacteria's fragile outer membrane and enter the cell to bind the DNA and RNA [33]. STM (WT), *ΔompA,* and *ΔompA*: pQE60-*ompA* were incubated with increasing concentrations of ceftazidime (0.25, 0.5, and 1 μg/ mL) **(Figure 5A)** and meropenem (0.01, 0.03, and 0.06 μg/ mL) **(Figure 5B)** and treated with propidium iodide at end of the incubation period to quantify the PI-positive dead bacterial population by flow cytometry. It was observed that in all three concentrations of ceftazidime **(Figure 5A.III, 5A.IV, 5A.V, and 5A.VI)** and meropenem **(Figure 5B.III, 5B.IV, 5B.V, and 5B.VI)** mentioned above, a significantly greater percentage of STM *ΔompA* (ceftazidime- **Figure 5A.III**- 80.76%, **5A.IV**- 83.34%, **5A.V**-82.31%, & **5A.VI**- cumulative trend and meropenem- **Figure 5B.III**- 28.31%, **5B.IV** - 38.55%, and **5B.V**- 42.66%, & **5B.VI-**cumulative trend) takes up propidium iodide compared to the STM (WT) (ceftazidime- **Figure 5A.III**- 66.72%, **5A.IV**- 66.99%, **5A.V**- 76.16%, & **5A.VI**- cumulative trend and meropenem- **Figure 5B.III**- 3.86%, **5B.IV**- 25.25%, and **5B.V**- 20.26%, & **5B.VI**- cumulative trend) and *ΔompA*: pQE60-*ompA* (ceftazidime- **Figure 5A.III**- 45.61%, **5A.IV**- 72.8%, **5A.V**- 73.91%, & **5A.VI**- cumulative trend and meropenem- **Figure 5A.III**- 5.22%, **5B.IV**- 25.75%, and **5B.V**- 32.92%, & **5B.VI**- cumulative trend), suggesting an enhanced killing of STM *ΔompA* by ceftazidime and meropenem-induced outer membrane depolarization. The significant increase in the PI-positive percent population of STM *ΔompA* affirmed the protective role of OmpA against the antibiotic-driven membrane disruption in *Salmonella* Typhimurium. Earlier, our data revealed that administration of β-lactam antibiotics against the wild type and OmpA mutant *Salmonella* Typhimurium produced an equivalent amount of ROS. Compared to STM *ΔompA,* the reduced killing of STM (WT) and *ΔompA*: pQE60-*ompA* strongly proved that the presence of OmpA helps the bacteria to fight against oxidative stress by maintaining the stability of the bacterial outer membrane during antibiotic stress.

### The administration of ceftazidime cleared bacterial infection from C57BL/6 mice more efficiently than meropenem

Besides antimicrobial resistance, bacterial cells have developed alternative strategies to survive antibiotic stress. Prolonged treatment of antibiotics to the infected hosts can generate antibiotic persisters which, constitute the transiently drug-tolerant phenotypic variants within isogenic populations. However, they do not proliferate in the presence of antibiotics, much in contrary to antibiotic-resistant bacteria [34, 35]. To date, the persisters are generally perceived as non-growing or slow-growing cells, and the reduced activity of antibiotic targets provides for their antibiotic tolerance [36, 37]. The bacteria can induce persistence and become resistant to antibiotic therapy by depolarizing the membrane potential [38]. However, the impact of outer membrane depolarization in the persistence of *Salmonella* Typhimurium is yet to be investigated. Our study demonstrated that the deletion of OmpA resulted in a significant increase in the outer membrane depolarization of STM *ΔompA* compared to STM (WT) in the presence of ceftazidime (0.25 μg/ml, 0.5 μg/ml, 1 μg/ml) and meropenem (0.01 μg/ml, 0.03 μg/ml, 0.06 μg/ml). Hence, we hypothesized that the increased membrane depolarization of STM *ΔompA* will help in developing antibiotic persistence. Much in contrary, we found that deletion of OmpA resulted in lesser percent viability of persisters as compared to wild-type in both planktonic **(Figure 6A)** as well as in biofilm **(Figure 6C)** culture following prolonged ceftazidime (50 μg/ml- 50X of MIC) treatment. However, no significant difference was reflected in the percent viability of STM (WT) and STM *ΔompA* in both planktonic **(Figure 6B)** and biofilm **(Figure 6D)** cultures following exposure to meropenem (50 μg/ml- 25X of MIC). We hypothesized that during ceftazidime treatment, the greater reduction in the persister population of OmpA deficient *Salmonella* might lead to better clearance of the bacteria from *in vivo* infection model. To verify this hypothesis, we have subjected 4 to 6 weeks old C57BL/6 mice infected with 10^6^ CFU of wild-type and mutant bacteria to the treatment of β lactam antibiotics **(Figure 6E-6G)**. Ceftazidime and meropenem (5 mg/ kg of body weight) were administered in the infected mice on the 2^nd^ and 4^th^-day post-infection by intraperitoneal injection **(Figure 6E)**. On the 5^th^-day post-infection, the mice were sacrificed, the liver and spleens were isolated, homogenized, and the organ lysate was plated to enumerate the bacterial load. In line with our expectation, it was found that, unlike meropenem, the administration of ceftazidime efficiently reduced the burden of STM *ΔompA* in the liver and spleen of C57BL/6 mice compared to the antibiotic-untreated mice **(Figure 6F and 6G)**.

**Figure 6.**
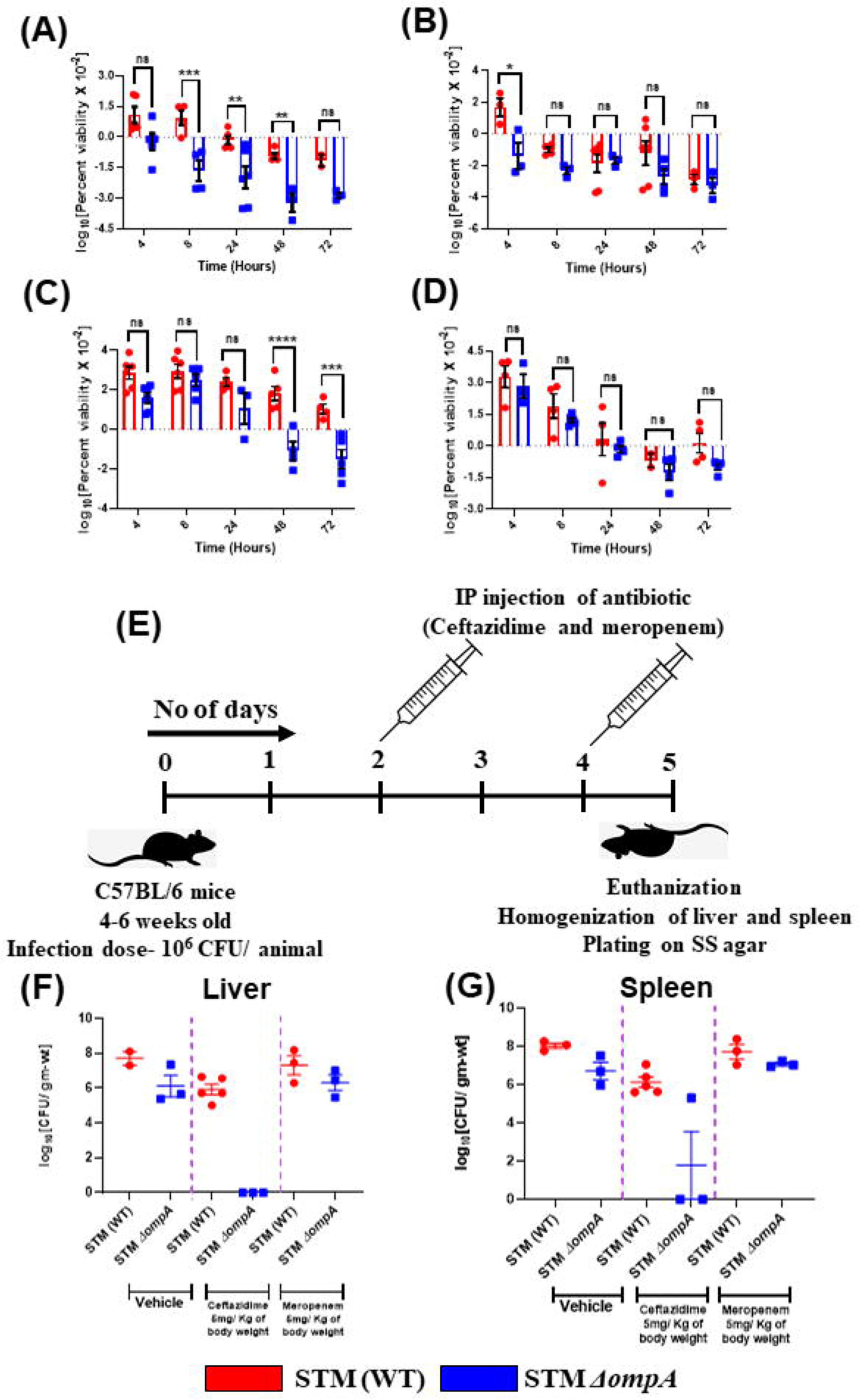
Organ burden of STM (WT) and *ΔompA* under the treatment of ceftazidime and meropenem (5 mg/ kg of body weight). (A-D) Calculating the antibiotic tolerant persister fraction of STM (WT) and *ΔompA* growing in planktonic culture (A-B) and bioflim (C-D) in the presence of ceftazidime (A and C) and meropenem (B and D) for 72 hours (n=2, N=3). (E-G) 4–6-week-old C57BL/6 mice were infected with 10^6^ CFU of STM (WT) and STM *ΔompA* (n=5). The mice were treated with ceftazidime and meropenem (5 mg/ kg of body weight) on the specified days (E). On the 5^th^ day post infection, the mice were sacrificed. The liver (F) and spleen (G) were collected and homogenised. The cell lysates were plated to enumerate the load of bacteria in each organ. The CFU obtained from the liver and spleen were normalised with the weight of the individual organs. ***(P)* *< 0.05, *(P)* **< 0.005, *(P)* ***< 0.0005, *(P)* ****< 0.0001, ns= non-significant, (2way ANOVA)**.

## 5.4 Discussion

The rapid emergence of drug resistance phenotype in non-typhoidal serovars of *Salmonella* is a significant obstacle in curing *Salmonella*-induced foodborne illness with antibiotic therapy. Invasive non-typhoidal *Salmonella* serovars, such as *S.* Typhimurium ST313, are responsible for bloodstream infection amongst the malnourished children and adults of sub-Saharan Africa [5]. Recent studies have revealed the appearance of XDR *S.* Typhimurium ST313 in Africa, possessing MDR, extended-spectrum β-lactamase, and azithromycin resistance, which has posed a significant threat to global health [4]. The evolution of the pathogen due to genome degradation has been assumed to be the primary reason behind the generation of antibiotic-resistant phenotypes [39]. The emergence of the MDR phenotype in another non-typhoidal *Salmonella* serovar *Salmonella* Typhimurium DT104 has also been reported. The infection in humans and cattle caused by this pathogen is mediated by *Salmonella* Genomic Island-1 (SGI-1), which confers protection against a wide range of antibiotics, encompassing ampicillin (*pse-1*), chloramphenicol/florfenicol (*floR*), streptomycin/ spectinomycin (*aadA2*), sulfonamides (*sul1*), and tetracycline (*tetG*) (ACSSuT) [40, 41]. Tigecycline and carbapenem are the latest anti-*Salmonella* drugs used to treat MDR and XDR typhoid fever. However, the continuous adaptation of the pathogen creates a potential risk of developing resistance against recommended antibiotics in the future, which further provides an opportunity to study new drugs and their potential target in detail. Gram-negative bacterial pathogens like *Escherichia coli, Acinetobacter baumannii, Klebsiella pneumoniae, Pseudomonas aeruginosa,* etc., use outer membrane-bound porins for various purposes, starting from maintaining the outer membrane stability to developing antibiotic resistance by regulating the permeability [9, 12, 13, 15, 16, 42–46].

In the current study, we have tried to delineate the contributions of the most abundant outer membrane porins of *Salmonella* Typhimurium such as OmpA, OmpC, OmpD, and OmpF in developing resistance against two β-lactam antibiotics, namely ceftazidime and meropenem. Ceftazidime and meropenem inhibit the biosynthesis of bacterial cell walls after binding to the penicillin-binding proteins [19, 47]. We have exposed the *ompA*, *ompC*, *ompD*, and *ompF* knockout strains of *Salmonella* to increasing concentrations of these drugs. It was found that the MIC of both the antibiotics reduces for the *ompA* knockout *Salmonella* compared to the wild-type and other porins (*ompC, ompD,* and *ompF*) knockout strains, suggesting that *Salmonella* Typhimurium OmpA plays an essential role to protect the pathogen from the β-lactam antibiotics. Despite the presence of the antibiotic, the partial reversal of the growth inhibition phenotype in *ompA* complemented strain further supported our conclusion. *Salmonella* Typhimurium OmpA has a unique structure. It has an outer membrane-bound β barrel subunit, which has four externally exposed extracellular loops and a periplasmic subunit that interacts with the peptidoglycan layer [17]. Compared to other porins, namely OmpC, OmpD, and OmpF, the smaller pore size of OmpA might be associated with slowing down the entry of antibiotic molecules across the bacterial outer membrane and protecting the bacteria. As the β-lactam antibiotics inhibit the biosynthesis of the bacterial cell walls, we hypothesized that in the absence of OmpA, these antibiotics would cause extensive damage to the bacterial external envelope. We measured the membrane disruption of the bacteria using a dye named DiBAC_4_, which enters the cell only when the cytosol has a higher positive charge due to outer membrane depolarization. The damaged outer membrane and cell wall facilitate the inflow of cations and reduce the cytosol's negative charge density, making it accessible towards DiBAC_4_. We found a negligible depolarization of the bacterial outer membrane when the β-lactam antibiotics were absent in the media, suggesting that the deletion of *ompA* itself is not lethal to the bacteria. The uninterrupted planktonic growth of STM (WT) and *ΔompA* in LB (data not shown) and MH broth (in the MIC determination experiment) proved our conviction to be true. We have also found a steady rise in membrane depolarization of all three bacterial strains with increased antibiotic concentrations. However, the membrane depolarization of STM *ΔompA* was significantly higher than the wild-type, and OmpA complemented strains in all three concentrations of antibiotics.

We further wanted to delineate the role of externally exposed extracellular loops of OmpA in antimicrobial resistance. In our previous study, we have proved that introducing mutations to the extracellular loops of OmpA can’t reduce the stability of the *Salmonella* outer membrane [27]. Our current study revealed that contrary to STM *ΔompA*, the OmpA loop mutants could resist antibiotic-mediated membrane depolarization and subsequent growth inhibition of the bacteria like wild-type and complemented strains. This result led us to assume that the fully folded, membrane-embedded, and functionally active structure of OmpA is required to maintain the outer membrane stability of *Salmonella* during antibiotic stress. We further speculated that the higher membrane depolarization of STM *ΔompA* could give rise to enhanced uptake of antibiotics, followed by severe damage of the bacterial morphology. Our experiments revealed that STM *ΔompA* takes up more antibiotic from the media than the wild-type and the complemented strains. Our confocal microscopy and atomic force microscopy results strongly supported the severe damage of the bacterial membrane in the absence of OmpA during antibiotics treatment. Irrespective of the target specificity, the bactericidal antibiotics produce reactive oxygen species to kill the bacterial pathogens [48]. To test this hypothesis, we quantified the external ROS generated by treating the wild type and *ompA* knockout *Salmonella* with a sub-lethal concentration of both antibiotics by DCFDA staining. The incubation of bacterial strains with antibiotics produced a comparable amount of ROS between the wild-type and the mutant *Salmonella*. However, we concluded that the STM *ΔompA* strain could not tolerate the oxidative stress generated by antibiotic treatment, unlike the STM (WT), which is the most probable reason behind its remarkable growth inhibition phenotype in response to the antibiotics.

Hence, we further looked into establishing a correlation between the bacterial outer membrane damage and viability in the presence and absence of antibiotics. We have stained the antibiotic-treated and untreated bacterial cells with propidium iodide to quantify the percentage of dead bacteria. Incubating STM *ΔompA* with increasing concentrations of β-lactam drugs increased the bacterial killing compared to the wild-type and complemented strains. Since previous studies in *Escherichia coli* and *Staphylococcus aureus* have demonstrated the association of membrane depolarization with the emergence of antibiotic persister population, we decided to extend the findings in the *Salmonella* Typhimurium model. Much in disparity, we found that the deletion of *ompA* from *Salmonella* resulted in decreased persister cell levels till 72h post-exposure to ceftazidime (50 μg/ml) and both in planktonic and biofilm cultures. Since deletion of *ompA* is also associated with increased cell death upon exposure to similar concentrations of ceftazidime (0.25 μg/mL, 0.5 μg/mL, 1 μg/mL), bacterial cell death might be overriding the antibiotic persistence. To support our *in vitro* observation, we further focused on verifying the ability of β-lactam drugs in clearing the bacterial infection in *in vivo* infection model. 4-6 weeks old C56BL/6 mice were infected with wild-type and *ompA* deficient *Salmonella*. The antibiotic treatment was started on the 2nd-day post-infection for the disease manifestation. Our data depicted that the administration of ceftazidime can reduce the burden of STM *ΔompA* compared to antibiotic-untreated mice, suggesting that *Salmonella* Typhimurium OmpA plays a crucial role in protecting the bacteria from β-lactam antibiotics.

To the best of our knowledge, we are reporting for the first time that apart from maintaining the stability of the bacterial outer membrane, OmpA directly takes part in antimicrobial resistance in *Salmonella* Typhimurium. The other major outer membrane porins (OmpC, OmpD, and OmpF) of *Salmonella* with larger pore sizes, don’t have any significant role in developing resistance against β-lactam drugs. In the absence of OmpA, bacteria consume more β-lactam drugs causing extensive damage to the bacterial outer membrane and making the bacteria highly susceptible to antibiotic-mediated oxidative stress.

## Materials and methods

### Bacterial strains, media, and culture conditions

The wild-type (WT) bacteria, *Salmonella enterica* serovar Typhimurium strain 14028S used in this study was a generous gift from Professor Michael Hensel, Department of Microbiology, University of Osnabruck, Germany. All the bacterial strains used in this study were revived from glycerol stock (stored in −80°C) and plated either only on LB agar (purchased from HiMedia) (for the wild-type *Salmonella*) or LB agar along with appropriate antibiotics like-kanamycin (50 μg/mL- for the *ompA* knockout strains), chloramphenicol (25 μg/mL- for the *ompC*, *ompD*, and *ompF* knockout strains), and ampicillin (50 μg/mL- for the complemented and the OmpA loop mutant strain). The complete list of bacterial strains used in this study has been listed below. (Description in Table 1). For all the experiments, a single bacterial colony from the LB agar plates (with or without antibiotics) was inoculated into the LB broth, followed by overnight incubation. The overnight-grown stationary phase culture was further subcultured at a 1: 100 ratio in a fresh LB tube and allowed to grow for 6 hours so that the bacteria attain the log phase. The OD of the bacterial cells was normalized to 0.1, which corresponds to 10^6^ CFU of bacteria. This normalized culture was used for all the experiments mentioned below.

**Table 1.** Strains used in this study.

| Strains/ plasmids | Characteristics | Source/ references |
| --- | --- | --- |
| *Salmonella enterica* serovar Typhimurium ATCC strain 14028S | Wild type (WT) | Gifted by Prof. M. Hensel |
| *S. Typhimurium* $\Delta ompA$ | Kan<sup>R</sup> | Laboratory stock |
| *S. Typhimurium* $\Delta ompA$: pQE60-*ompA* | Kan<sup>R</sup>, Amp<sup>R</sup> | Laboratory stock |
| *S. Typhimurium* $\Delta ompA$: pQE60 | Kan<sup>R</sup>, Amp<sup>R</sup> | Laboratory stock |
| *S. Typhimurium* $\Delta ompC$ | Chl<sup>R</sup> | Laboratory stock |
| *S. Typhimurium* $\Delta ompD$ | Kan<sup>R</sup> | Laboratory stock |
| *S. Typhimurium* $\Delta ompF$ | Chl<sup>R</sup> | Laboratory stock |
| *S. Typhimurium* $\Delta ompA$: pQE60-*ompA* L1-1 | Kan<sup>R</sup>, Amp<sup>R</sup> | Laboratory stock |
| *S. Typhimurium* $\Delta ompA$: pQE60-*ompA* L1-2 | Kan<sup>R</sup>, Amp<sup>R</sup> | Laboratory stock |
| *S. Typhimurium* $\Delta ompA$: pQE60-*ompA* L2-1 | Kan<sup>R</sup>, Amp<sup>R</sup> | Laboratory stock |
| *S. Typhimurium* $\Delta ompA$:<br>pQE60-*ompA* L2-2 | Kan<sup>R</sup>, Amp<sup>R</sup> | Laboratory stock |
| *S. Typhimurium* $\Delta ompA$:<br>pQE60-*ompA* L3-1 | Kan<sup>R</sup>, Amp<sup>R</sup> | Laboratory stock |
| *S. Typhimurium* $\Delta ompA$:<br>pQE60-*ompA* L4-1 | Kan<sup>R</sup>, Amp<sup>R</sup> | Laboratory stock |

### Determination of the minimal inhibitory concentration (MIC) of ceftazidime and meropenem

The log phase cultures of STM (WT), *ΔompA*, *ΔompA*:pQE60-*ompA*, *ΔompC*, *ΔompD*, *ΔompF*, *ΔompA*:pQE60-*ompA*-L1-1, *ΔompA*:pQE60-*ompA*-L1-2, *ΔompA*:pQE60-*ompA*-L2-1, *ΔompA*:pQE60-*ompA*-L2-2, *ΔompA*:pQE60-*ompA*-L3-1, and *ΔompA*:pQE60-*ompA*-L4-1 (OD adjusted to 0.1) were used to determine the MIC of ceftazidime and meropenem. A working stock of ceftazidime (concentration- 256 μg/ mL) was prepared in cation-adjusted Muller-Hinton broth and serially diluted in the wells of a 96-well plate with freshly prepared (autoclaved) Muller-Hinton broth to produce 100 μL volume of the following concentrations 256-, 128-, 64-, 32-, 16-, 8-, 4-, 2-, 1-, 0.5-, and 0 μg/ mL, respectively. 100 μL of bacterial cultures where the OD was normalized to 0.1, were added in each well of the 96-well plate, which made the final volume of the media 200 μL per well, with ceftazidime concentrations as follows 128-, 64-, 32-, 16-, 8-, 4-, 2-, 1-, 0.5-, 0.25-, and 0 μg/ mL, respectively. For meropenem, the final concentrations in each well were kept as follows 8-, 4-, 2-, 1-, 0.5-, 0.25-, 0.125-, 0.06-, 0.03-, 0.01-, and 0 μg/ mL, respectively. Two wells-one with antibiotic and the other without any antibiotic, were kept in the same plate without any inoculation as blanks. The plates were incubated for 18 to 24 hours in a shaking incubator at 37°C temperature at 170 rpm. At the end of the incubation period, the plates were subjected to OD measurement by a TECAN microplate reader to investigate the bacterial growth inhibition and MIC determination.

### Determination of bacterial viability by resazurin assay

The log phase cultures of STM (WT), *ΔompA*, *ΔompC*, *ΔompD*, *ΔompF*, *ΔompA*:pQE60-*ompA*-L1-1, *ΔompA*:pQE60-*ompA*-L1-2, *ΔompA*:pQE60-*ompA*-L2-1, *ΔompA*:pQE60-*ompA*-L2-2, *ΔompA*:pQE60-*ompA*-L3-1, and *ΔompA*:pQE60-*ompA*-L4-1 (OD adjusted to 0.1) were used to determine the viability in the presence or absence of ceftazidime and meropenem. The bacterial cells were subjected to increasing concentrations of antibiotic treatment (protocol mentioned above) for 16 to 18 hours. At the end of the incubation period, 20 μL of resazurin solution from a stock of 0.2 mg/ mL was added to the bacterial suspensions present in the wells of 96-well plate and further incubated for 2 hours in dark conditions. The fluorescence intensity of resazurin (excitation-540 nm and emission-590 nm) was measured with the help of a TECAN microplate reader. The fluorescence intensity obtained from the well without any antibiotic was considered a hundred percent viable, and the percent viability for the antibiotic-treated samples was calculated.

### Determination of the ROS generation upon antibiotic treatment

The log phase cultures of STM (WT) and *ΔompA* were used to determine ROS generation in the presence or absence of ceftazidime and meropenem. The bacterial cells (prepared according to the protocol mentioned above) were incubated with sub-lethal concentrations of ceftazidime (0.25 μg/ mL) and meropenem (0.01 μg/ mL) for 18 to 24 hours. At the end of the incubation period, the cells were treated with DCFDA (10 μM) for 15 minutes and washed with sterile PBS once. The washed cells were immediately subjected to analysis by flow cytometry (BD FACSVerse by BD Biosciences-US) using 492 excitation and 517 emission channels, respectively. The results were analyzed by BD FACSuite software.

### Measurement of outer membrane depolarization

The depolarization of bacterial outer membrane upon antibiotic treatment was measured using a dye called DiBAC_4_. The log phase cultures of STM (WT), *ΔompA*, *ΔompA*:pQE60-*ompA*, *ΔompA*:pQE60-*ompA*-L1-1, *ΔompA*:pQE60-*ompA*-L1-2, *ΔompA*:pQE60-*ompA*-L2-1, *ΔompA*:pQE60-*ompA*-L2-2, *ΔompA*:pQE60-*ompA*-L3-1, and *ΔompA*:pQE60-*ompA*-L4-1 were prepared according to the aforementioned protocol. The bacterial cells were incubated with increasing concentrations of ceftazidime and meropenem for 18 to 24 hours. At the end of the incubation period, the cells were treated with DiBAC_4_ (1 μg/ mL) for 15 minutes and washed with sterile PBS once. The washed cells were immediately subjected to analysis by flow cytometry (BD FACSVerse by BD Biosciences-US), and the results were analyzed by BD FACSuite software.

### Measurement of the dead bacterial population

Propidium iodide (working concentration-1μg/ mL) was used to estimate the bacterial death upon antibiotic treatment. Propidium iodide is an intercalating dye that enters the dead bacterial cells and binds to the bases of DNA. The propidium iodide bound to the DNA starts fluorescing. The bacterial cells were incubated with increasing concentrations of meropenem for 18 to 24 hours. At the end of the incubation period, the cells were treated with propidium iodide for 15 minutes and washed with sterile PBS once. The washed cells were immediately subjected to analysis by flow cytometry (BD FACSVerse by BD Biosciences-US), and the results were analyzed by BD FACSuite software.

### Measuring the entry of meropenem by HPLC

HPLC from Agilent Technologies (1120 Compact LC) with a C18 column as a stationary phase (at 30°C) was used to quantify the meropenem uptake by the bacteria. A mixture of 0.1% aqueous acetic acid (solution A) and methanol (solution B) was used as a mobile phase with a gradient program (Description in Table 2). The flow rate of the mobile phase was kept at 1 mL/ minute for chromatographic separation. The detection wavelength for meropenem was fixed at 300 nm. The elution of meropenem happened in between 13-14 minutes. The standard curve was formulated with known concentrations of meropenem (5, 25, 50, 75, and 100 μg/ mL). The Muller-Hinton broth without any antibiotic was used as a blank. The area under the peak was calculated to estimate the availability of meropenem and plotted against the antibiotic concentration to form the straight line. The exponentially growing cultures of STM (WT), *ΔompA*, and *ΔompA*:pQE60-*ompA* (~ 10^7^ CFU of bacteria) were subjected to the treatment with a very high dose of meropenem (concentration ~ 100-150 μg/ mL). One hour after incubation at 37°C, we have harvested the cells by centrifugation at 5000 rpm for 20 minutes. The culture supernatant was collected and filter-sterilized. 20 μL of this culture supernatant was used to quantify the remaining concentration of meropenem. The results obtained from the instrument were analyzed by EZChrome Elite software, and the area under the curve was measured for estimating the concentration of available antibiotics in the media.

**Table 2.** The gradient program of the mobile phase used for HPLC.

| **Time in minutes** | **Solution A**<br>**(0.1% aqueous acetic acid)** | **Solution B**<br>**(Methanol)** |
| --- | --- | --- |
| 0 | 95% | 5% |
| 2 | 95% | 5% |
| 18 | 75% | 25% |
| 20 | 75% | 25% |
| 25 | 50% | 50% |
| 28 | 50% | 50% |
| 29 | 95% | 5% |
| 35 | 95% | 5% |

### Assessing the damage of the bacterial envelope (cell wall and cell membrane) by confocal LASER scanning microscopy (CLSM)

To study the outer membrane damage of the bacteria upon treatment with antibiotics, the wild-type, mutant, and complemented strains of *Salmonella* were subjected to increasing concentrations of meropenem. After 16 hours of incubation, the antibiotic-treated or untreated bacterial cells were fixed with 3.5% PFA. To stain the outer membrane of the bacteria, the fixed bacterial cells were incubated with FM 4-64 (excitation/ emission-514/640 nm) (0.01 μg/ mL) for 30 minutes. The DNA of the bacterial cells was visualized by DAPI staining (0.01 μg/ mL). The images were acquired and analyzed by confocal laser scanning microscopy 880 (ZEISS) and ZEN black software.

### Assessing the bacterial morphology by atomic force microscopy (AFM)

The exponential phase cultures of STM (WT), *ΔompA*, and *ΔompA*:pQE60-*ompA* were subjected to the treatment with a sublethal concentration of meropenem (0.03 μg/ mL). Antibiotic untreated cells were also allowed to grow under similar growth conditions. 16 to 18 hours post-infection; the cells were fixed with 3.5% PFA and washed with double autoclaved Milli-Q water. The washed cells were further diluted by four folds with Milli-Q water. A 100 μL of this diluted sample was dropcasted on sterile glass coverslips. The completely airdried sample on the coverslip was taken to NX10 Atomic Force Microscope (AFM) for image acquisition.

### Animal experiments

4 to 6 weeks old C57BL/6 mice were infected with 10^6^ CFU of wild type and *ΔompA*. For the manifestation of the disease, the mice were kept undisturbed for a day. On the 2^nd^-day post-infection, the infected mice were administered with ceftazidime and meropenem (5 mg/ kg of body weight) separately. The antibiotic treatment was provided on every alternative day. Antibiotic untreated mice were kept as controls. On the 5^th^-day post-infection, the mice were sacrificed, and the infected organs such as the liver and spleen were collected. The organs were homogenized with sterile glass beads, and the lysates were plated on *Salmonella Shigella* agar plates to enumerate the bacterial load. The CFU obtained after plating was normalized with the weight of the individual organs, and the log value of the normalized CFU was plotted.

### Statistical analysis

Each experiment has been independently repeated (with at least two biological replicates) multiple times (The n=technical replicates and N= biological replicates have been mentioned in the figure legends wherever applicable). The graphs and the statistics were formulated by GraphPad Prism 8.4.3 software with the help of the numerical data points obtained from different experiments. For multiple comparisons (data points obtained from the growth inhibition experiment, percent viability calculation for the porin knockout strains, cumulative trend of membrane depolarization, and bacterial death), 2wayANOVA was used. To quantify the HPLC data, the estimation of percent viability for the OmpA loop mutants and the calculation of the persister population unpaired students t-test were used. In all the cases, the *p* values below 0.05 were considered significant. The results are expressed as mean ± SEM. Differences between experimental groups were deemed to be significant for *p*< 0.05.

## Abbreviations

STM: *Salmonella Typhimurium*
OmpA: Outer membrane protein A
OmpC: Outer membrane protein C
OmpD: Outer membrane protein D
OmpF: Outer membrane protein F
MHB: Muller-Hinton broth
PFA: Para-formaldehyde
FM4-64: N-(3-Triethylammoniumpropyl)-4-(6-(4-(Diethylamino) Phenyl) Hexatrienyl) Pyridinium Dibromide
DAPI: 4′,6-diamidino-2-phenylindole
ROS: Reactive oxygen species
DiBAC_4_: Bis-(1,3-Dibutylbarbituric Acid)Trimethine Oxonol
HPLC: High performance liquid chromatography

## Author Contributions

ARC and DM equally contributed to the construction of the manuscript. ARC and DC conceived the study and designed the experiments. ARC and DM performed all the experiments, participated in the acquisition and analysis of the data. AS performed the experiments with ARC and DM. ARC constructed the figures and wrote the original draft of the manuscript. DM, AS, and DC participated in the proofreading and editing of the manuscript. DC supervised the study. All the authors have read and approved the manuscript.

## Acknowledgments

We sincerely thank the departmental confocal facility, departmental real-time PCR facility, central bioimaging facility, central flow cytometry facility at IISc. Mr. Puneeth and Ms. Navya from the departmental confocal facility are acknowledged for their assistance in image acquisition. Ms. Leepika and Ms. Sharon from the central flowcytometry facility are duly acknowledged for their help in flow cytometry data acquisition. We sincerely acknowledge the assistance provided by the Central Animal Facility of IISc. Dr. Shivjee Sah from Professor Umesh Varshney’s laboratory MCB, IISc, is duly thanked for constructing the *ompA*-pQE60 recombinant plasmid.

## Availability of data and materials

All data generated and analyzed during this study, including the supplementary information files, have been incorporated in this article. The data is available from the corresponding author on reasonable request.

## Declarations

### Ethics statement

The Institutional Animal Ethics Committee approved all the animal experiments, and the Guidelines provided by National Animal Care were strictly followed during animal experiments. (Registration No: 48/1999/CPCSEA).

### Consent for publication

Not applicable.

## Supplementary Figures

**Figure S1.**
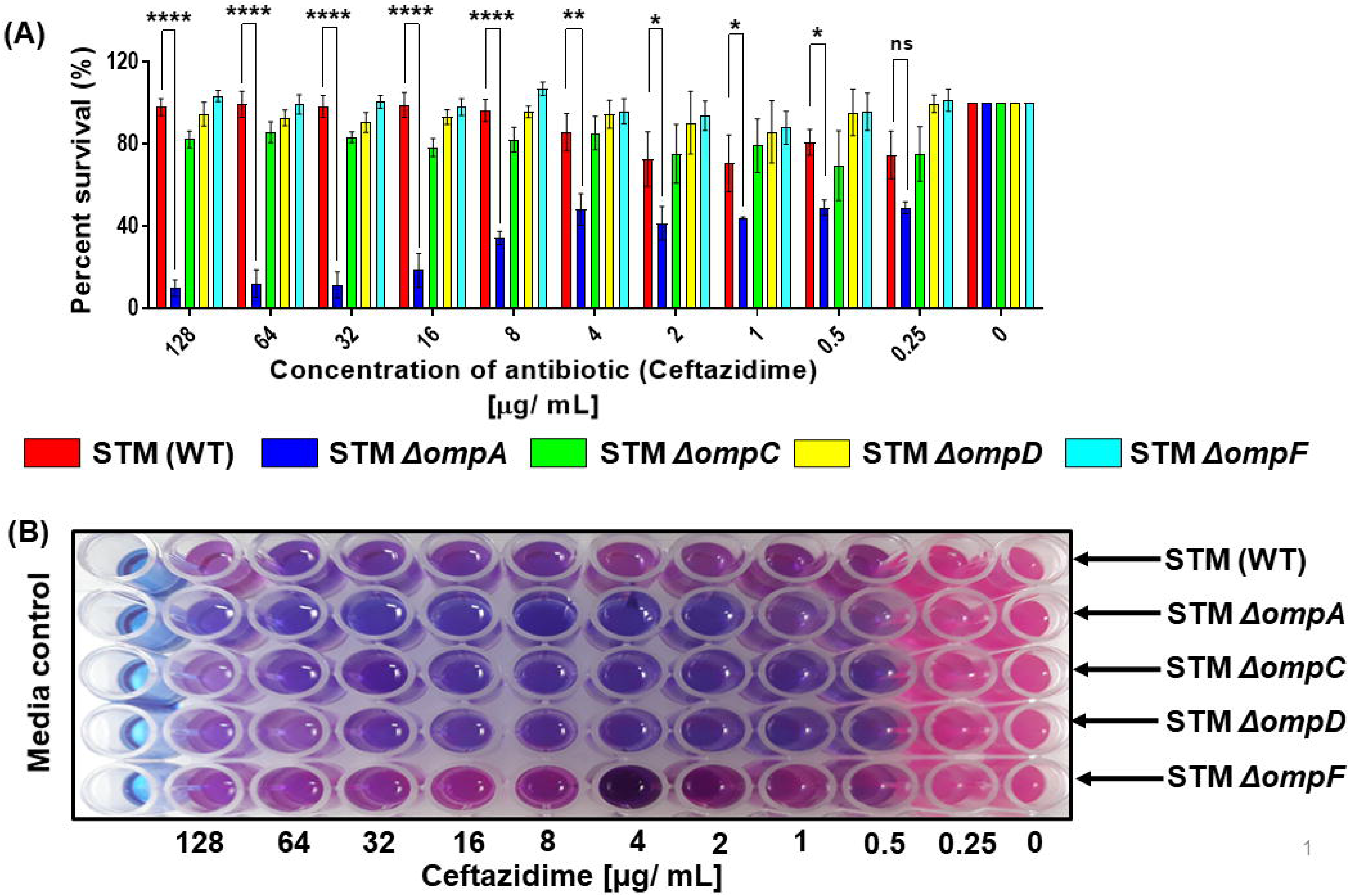
Estimating the percent viability of bacteria in the presence of ceftazidime by resazurin assay. (A) Determination of the percent viability of STM (WT), *ΔompA*, *ΔompC*, *ΔompD*, and *ΔompF* growing in cation-adjusted Mueller-Hinton broth in the presence of ceftazidime by resazurin assay (N=3). (B) The pictorial representation of antibiotic treated or untreated STM (WT), *ΔompA*, *ΔompC*, *ΔompD*, and *ΔompF* in the rpesensence of resazurin. ***(P)* *< 0.05, *(P)* **< 0.005, *(P)* ***< 0.0005, *(P)* ****< 0.0001, ns= non-significant, (2way ANOVA)**.

**Figure S2.**
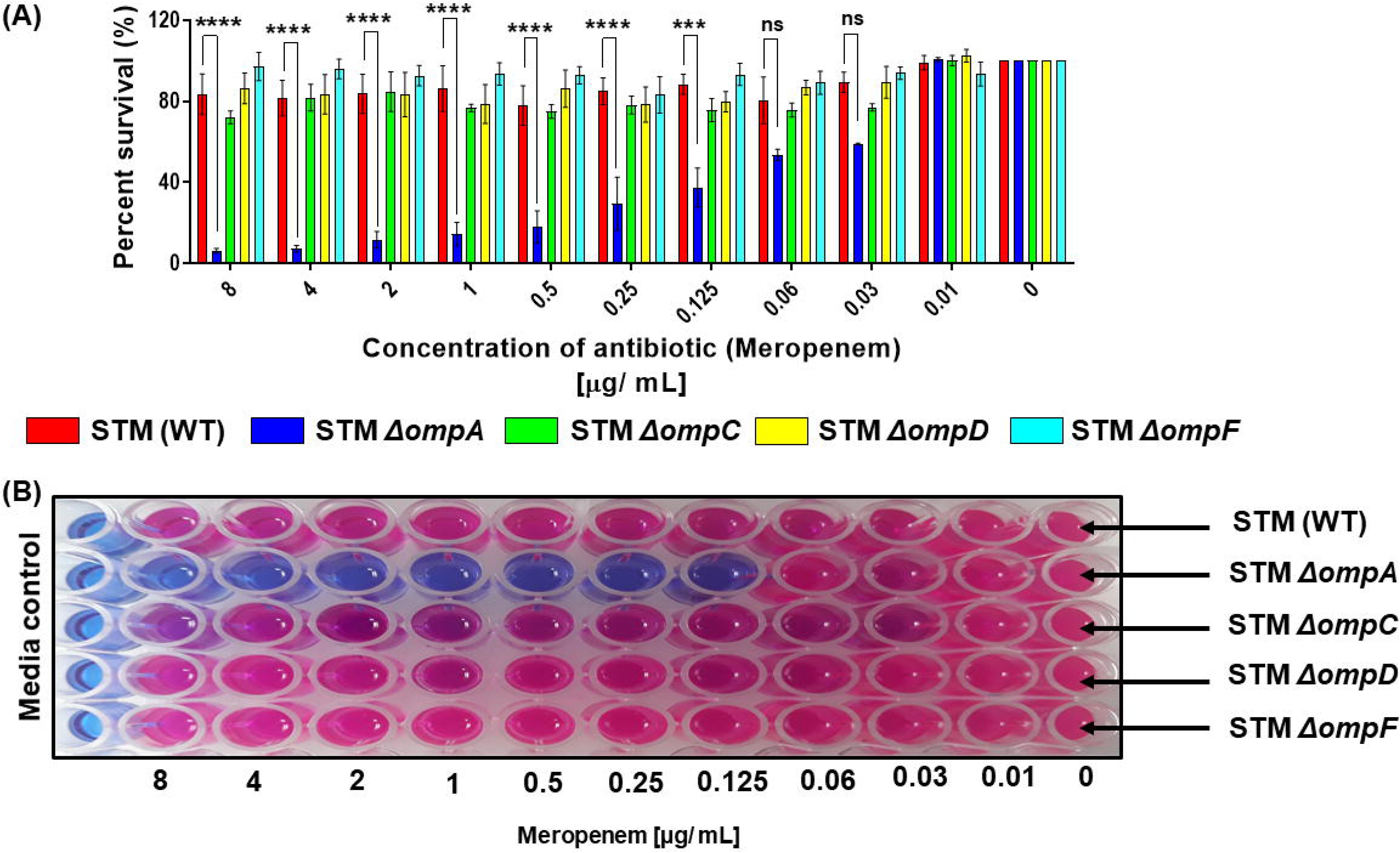
Estimating the percent viability of bacteria in the presence of meropenem by resazurin assay. (A) Determination of the percent viability of STM (WT), *ΔompA*, *ΔompC*, *ΔompD*, and *ΔompF* growing in cation-adjusted Mueller-Hinton broth in the presence of meropenem by resazurin assay (N=3). (B) The pictorial representation of antibiotic treated or untreated STM (WT), *ΔompA*, *ΔompC*, *ΔompD*, and *ΔompF* in the rpesensence of resazurin. ***(P)* *< 0.05, *(P)* **< 0.005, *(P)* ***< 0.0005, *(P)* ****< 0.0001, ns= non-significant, (2way ANOVA)**.

**Figure S3.**
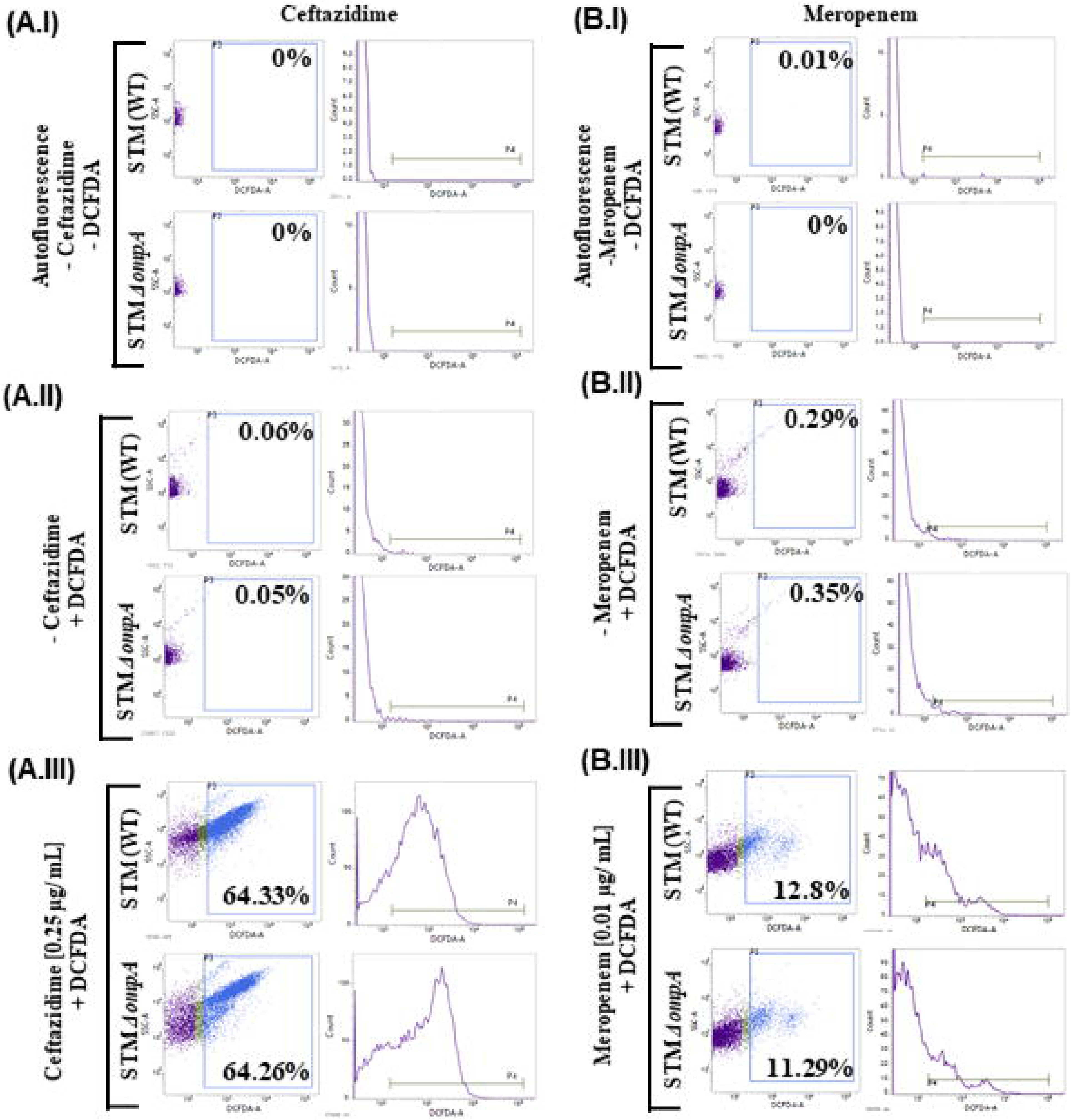
Exposure of the wild-type and the *ompA* knockout strains of *Salmonella* Typhimurium to the sublethal concentration of β-lactam antibiotics resulted in the generation of comparable amount of ROS. Studying the generation of ROS in STM (WT) and *ΔompA*, growing in cation-adjusted Mueller-Hinton broth in the presence of sublethal concentrations of β lactam antibiotics - (A.I-A.III) ceftazidime (0.25 μg/ mL) and (B.I- B.III) meropenem (0.01 μg/ mL) by DCFDA staining by flow cytometry (N=2). The final concentration of DCFDA used to measure the ROS burden of bacteria was 10 μM. The representative image corresponds to one single experiment of two independently done experiments. **The dot plot (SSC-A Vs. DCFDA-A) and the histogram (Count Vs. DCFDA-A) have been obtained from BD FACSuite software.**

**Figure S4.**
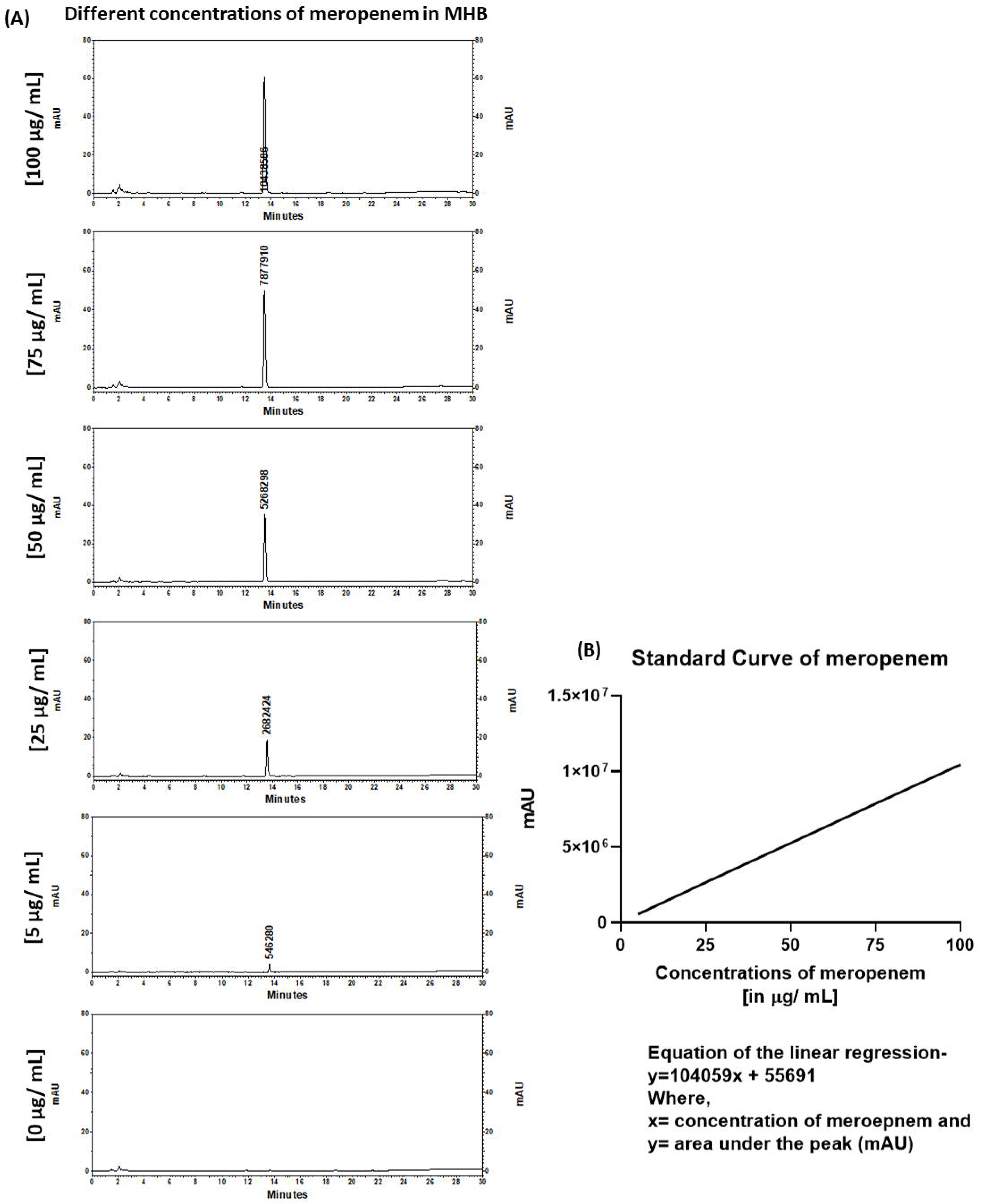
Generation of the standard curve with the known concentrations of meropenem in Muller-Hinton broth. The standard curve was constructed with known concentrations of meropenem (0, 5, 25, 50, 75, and 100 μg/ mL) in the Muller-Hinton broth. (A) The area under the peak was calculated to estimate the availability of the antibiotic and (B) plotted against the antibiotic concentration to form the straight line.

## References

1. Wang, X., et al., Antibiotic Resistance in Salmonella Typhimurium Isolates Recovered From the Food Chain Through National Antimicrobial Resistance Monitoring System Between 1996 and 2016. Front Microbiol, 2019. 10: p. 985.

2. Kurtz, J.R., J.A. Goggins, and J.B. McLachlan, Salmonella infection: Interplay between the bacteria and host immune system. Immunol Lett, 2017. 190: p. 42–50.

3. Branchu, P., M. Bawn, and R.A. Kingsley, Genome Variation and Molecular Epidemiology of Salmonella enterica Serovar Typhimurium Pathovariants. Infect Immun, 2018. 86(8).

4. Van Puyvelde, S., et al., An African Salmonella Typhimurium ST313 sublineage with extensive drug-resistance and signatures of host adaptation. Nat Commun, 2019. 10(1): p. 4280.

5. Collaborators, G.B.D.N.-T.S.I.D., The global burden of non-typhoidal salmonella invasive disease: a systematic analysis for the Global Burden of Disease Study 2017. Lancet Infect Dis, 2019. 19(12): p. 1312–1324.

6. Amuasi, J.H. and J. May, Non-typhoidal salmonella: invasive, lethal, and on the loose. Lancet Infect Dis, 2019. 19(12): p. 1267–1269.

7. Feasey, N.A., et al., Invasive non-typhoidal salmonella disease: an emerging and neglected tropical disease in Africa. Lancet, 2012. 379(9835): p. 2489–2499.

8. Leekitcharoenphon, P., et al., Global Genomic Epidemiology of Salmonella enterica Serovar Typhimurium DT104. Appl Environ Microbiol, 2016. 82(8): p. 2516–26.

9. May, K.L. and M. Grabowicz, The bacterial outer membrane is an evolving antibiotic barrier. Proc Natl Acad Sci U S A, 2018. 115(36): p. 8852–8854.

10. Nikaido, H., Molecular basis of bacterial outer membrane permeability revisited. Microbiol Mol Biol Rev, 2003. 67(4): p. 593–656.

11. Pages, J.M., C.E. James, and M. Winterhalter, The porin and the permeating antibiotic: a selective diffusion barrier in Gram-negative bacteria. Nat Rev Microbiol, 2008. 6(12): p. 893–903.

12. Choi, U. and C.R. Lee, Distinct Roles of Outer Membrane Porins in Antibiotic Resistance and Membrane Integrity in Escherichia coli. Front Microbiol, 2019. 10: p. 953.

13. Smani, Y., et al., Role of OmpA in the multidrug resistance phenotype of Acinetobacter baumannii. Antimicrob Agents Chemother, 2014. 58(3): p. 1806–8.

14. Kwon, H.I., et al., Outer membrane protein A contributes to antimicrobial resistance of Acinetobacter baumannii through the OmpA-like domain. J Antimicrob Chemother, 2017. 72(11): p. 3012–3015.

15. Iyer, R., et al., Acinetobacter baumannii OmpA Is a Selective Antibiotic Permeant Porin. ACS Infect Dis, 2018. 4(3): p. 373–381.

16. Llobet, E., et al., Klebsiella pneumoniae OmpA confers resistance to antimicrobial peptides. Antimicrob Agents Chemother, 2009. 53(1): p. 298–302.

17. van der Heijden, J., et al., Salmonella Rapidly Regulates Membrane Permeability To Survive Oxidative Stress. mBio, 2016. 7(4).

18. Roy Chowdhury, A., et al., Salmonella Typhimurium outer membrane protein A (OmpA) renders protection against nitrosative stress by promoting SCV stability in murine macrophages. bioRxiv, 2021: p. 2021.02.12.430987.

19. Shirley, M., Ceftazidime-Avibactam: A Review in the Treatment of Serious Gram-Negative Bacterial Infections. Drugs, 2018. 78(6): p. 675–692.

20. Dhillon, S., Meropenem/Vaborbactam: A Review in Complicated Urinary Tract Infections. Drugs, 2018. 78(12): p. 1259–1270.

21. Howe, R.A., J.M. Andrews, and B.W.P.o.S. Testing, BSAC standardized disc susceptibility testing method (version 11). J Antimicrob Chemother, 2012. 67(12): p. 2783–4.

22. Epand, R.F., et al., Depolarization, bacterial membrane composition, and the antimicrobial action of ceragenins. Antimicrob Agents Chemother, 2010. 54(9): p. 3708–13.

23. Te Winkel, J.D., et al., Analysis of Antimicrobial-Triggered Membrane Depolarization Using Voltage Sensitive Dyes. Front Cell Dev Biol, 2016. 4: p. 29.

24. Han, F.F., et al., Comparing bacterial membrane interactions and antimicrobial activity of porcine lactoferricin-derived peptides. J Dairy Sci, 2013. 96(6): p. 3471–87.

25. Soren, O., et al., Antimicrobial Peptide Novicidin Synergizes with Rifampin, Ceftriaxone, and Ceftazidime against Antibiotic-Resistant Enterobacteriaceae In Vitro. Antimicrob Agents Chemother, 2015. 59(10): p. 6233–40.

26. Cheng, M., et al., Ramoplanin at bactericidal concentrations induces bacterial membrane depolarization in Staphylococcus aureus. Antimicrob Agents Chemother, 2014. 58(11): p. 6819–27.

27. Chowdhury, A.R., D. Hajra, and D. Chakravortty, The extracellular loops of Salmonella Typhimurium outer membrane protein A (OmpA) maintain the stability of Salmonella containing vacuole (SCV) in murine macrophages and protect the bacteria from autophagy-dependent lysosomal degradation. bioRxiv, 2021: p. 2021.11.07.467609.

28. Van Acker, H. and T. Coenye, The Role of Reactive Oxygen Species in Antibiotic-Mediated Killing of Bacteria. Trends Microbiol, 2017. 25(6): p. 456–466.

29. Cabiscol, E., J. Tamarit, and J. Ros, Oxidative stress in bacteria and protein damage by reactive oxygen species. Int Microbiol, 2000. 3(1): p. 3–8.

30. Zhao, X. and K. Drlica, Reactive oxygen species and the bacterial response to lethal stress. Curr Opin Microbiol, 2014. 21: p. 1–6.

31. Van Acker, H., et al., The Role of Reactive Oxygen Species in Antibiotic-Induced Cell Death in Burkholderia cepacia Complex Bacteria. PLoS One, 2016. 11(7): p. e0159837.

32. Dwyer, D.J., et al., Antibiotics induce redox-related physiological alterations as part of their lethality. Proc Natl Acad Sci U S A, 2014. 111(20): p. E2100–9.

33. Rosenberg, M., N.F. Azevedo, and A. Ivask, Propidium iodide staining underestimates viability of adherent bacterial cells. Sci Rep, 2019. 9(1): p. 6483.

34. Brauner, A., et al., Distinguishing between resistance, tolerance and persistence to antibiotic treatment. Nat Rev Microbiol, 2016. 14(5): p. 320–30.

35. Bartell, J.A., et al., Bacterial persisters in long-term infection: Emergence and fitness in a complex host environment. PLoS Pathog, 2020. 16(12): p. e1009112.

36. Balaban, N.Q., et al., Bacterial persistence as a phenotypic switch. Science, 2004. 305(5690): p. 1622–5.

37. Lewis, K., Persister cells. Annu Rev Microbiol, 2010. 64: p. 357–72.

38. Benarroch, J.M. and M. Asally, The Microbiologist's Guide to Membrane Potential Dynamics. Trends Microbiol, 2020. 28(4): p. 304–314.

39. Pulford, C.V., et al., Stepwise evolution of Salmonella Typhimurium ST313 causing bloodstream infection in Africa. Nat Microbiol, 2021. 6(3): p. 327–338.

40. Boyd, D., et al., Characterization of variant Salmonella genomic island 1 multidrug resistance regions from serovars Typhimurium DT104 and Agona. Antimicrob Agents Chemother, 2002. 46(6): p. 1714–22.

41. Poppe, C., et al., Salmonella typhimurium DT104: a virulent and drug-resistant pathogen. Can Vet J, 1998. 39(9): p. 559–65.

42. Delcour, A.H., Outer membrane permeability and antibiotic resistance. Biochim Biophys Acta, 2009. 1794(5): p. 808–16.

43. Park, J.S., et al., Mechanism of anchoring of OmpA protein to the cell wall peptidoglycan of the gram-negative bacterial outer membrane. FASEB J, 2012. 26(1): p. 219–28.

44. Nie, D., et al., Outer membrane protein A (OmpA) as a potential therapeutic target for Acinetobacter baumannii infection. J Biomed Sci, 2020. 27(1): p. 26.

45. Samanta, S., et al., Getting Drugs through Small Pores: Exploiting the Porins Pathway in Pseudomonas aeruginosa. ACS Infect Dis, 2018. 4(10): p. 1519–1528.

46. Krishnan, S. and N.V. Prasadarao, Outer membrane protein A and OprF: versatile roles in Gram-negative bacterial infections. FEBS J, 2012. 279(6): p. 919–31.

47. Yang, Y., N. Bhachech, and K. Bush, Biochemical comparison of imipenem, meropenem and biapenem: permeability, binding to penicillin-binding proteins, and stability to hydrolysis by beta-lactamases. J Antimicrob Chemother, 1995. 35(1): p. 75–84.

48. Dwyer, D.J., M.A. Kohanski, and J.J. Collins, Role of reactive oxygen species in antibiotic action and resistance. Curr Opin Microbiol, 2009. 12(5): p. 482–9.

